# Tagging active neurons by soma-targeted Cal-Light

**DOI:** 10.1101/2021.10.13.464095

**Authors:** Jung Ho Hyun, Kenichiro Nagahama, Ho Namkung, Neymi Mignocchi, Patrick Hannan, Sarah Krüssel, Chuljung Kwak, Abigail McElroy, Bian Liu, Mingguang Cui, Seunghwan Lee, Dongmin Lee, Richard L. Huganir, Akira Sawa, Hyung-Bae Kwon

**Affiliations:** Solomon H. Snyder Department of Neuroscience, Johns Hopkins University School of Medicine, Baltimore, MD 21205, USA; Max Planck Florida Institute for Neuroscience, Jupiter, Florida 33458, USA; Department of Psychiatry, Neuroscience, Biomedical Engineering, and Genetic Medicine, Johns Hopkins University School of Medicine, Baltimore, MD 21205, USA; Department of Anatomy, Korea University College of Medicine, Seoul, Republic of Korea; Department of Mental Health, Johns Hopkins University Bloomberg School of Public Health, Baltimore, MD 21025, USA; Department of Brain and Cognitive Sciences, DGIST, Daegu, Republic of Korea

## Abstract

Verifying causal effects of neural circuits is essential for proving direct a circuit-behavior relationship. However, techniques for tagging only active neurons with high spatiotemporal precision remain at the beginning stages. Here we developed the soma-targeted Cal-Light (ST-Cal-Light) which selectively converts somatic calcium rise triggered by action potentials into gene expression. Such modification simultaneously increases the signal-to-noise ratio (SNR) of reporter gene expression and reduces the light requirement for successful labeling. Because of the enhanced efficacy, the ST-Cal-Light enables the tagging of functionally engaged neurons in various forms of behaviors, including context-dependent fear conditioning, leverpressing choice behavior, and social interaction behaviors. We also targeted kainic acid-sensitive neuronal populations in the hippocampus which subsequently suppressed seizure symptoms, suggesting ST-Cal-Light’s applicability in controlling disease-related neurons. Furthermore, the generation of a conditional ST-Cal-Light knock-in (KI) mouse provides an opportunity to tag active neurons in a region- or cell-type specific manner via crossing with other Cre-driver lines. Thus, the versatile ST-Cal-Light system links somatic action potentials to behaviors with high temporal precision, and ultimately allows functional circuit dissection at a single cell resolution.

## Introduction

Selective labeling and manipulation of behaviorally-engaged neuronal populations are critical for verifying their causal functions. To target active neuronal populations, immediate early gene (IEG)-based tagging systems have been widely used^1–5^. However, IEGs and their derivatives have limited behavioral applicability because of their poor temporal resolution (several hours) and weak coupling to cell firing^6^. To overcome these limitations, recently developed techniques implemented a dual light- and calcium-dependent switch system allowing investigators to label active cells with higher temporal precision^7–10^. In brief, these systems used “AND gate” logic, such that gene expression is initiated when both Ca^2+^ and light are present. Because action potentials causes opening of voltage-dependent Ca^2+^ channels followed by Ca^2+^ influx, light illumination during a specific time period enables gene expression only in neurons active during the behavior.

We previously reported that the Cal-Light technique was useful for specific labeling of the neuronal populations engaged with a bout of behavior during a lever pressing task in mice: inhibiting activity of the labeled neurons resulted in behavioral impairment^8^. Labeling and controlling irrelevant population of neurons that are not associated with lever pressing did not cause deficits in lever pressing behavior, indicating Cal-Light’s high selectivity^8^. Recently, the Cal-Light technique was proven to be useful for exclusively tagging and manipulating neural circuits underlying high cognitive behaviors, demonstrating its broad applicability^11^.

Despite such significant improvement, some caveats still remain. One major weakness could be a portion of gene expression caused by spontaneously occurring background Ca^2+^ signals. Such transient Ca^2+^ influxes arising through various routes would result in accumulation of action potential-independent labeling. For instance, Ca^2+^ concentration increases are caused not only by somatic action potentials, but also by local dendritic Ca^2+^ spikes or synaptic activity in dendrites or dendritic spines^12–14^ Internal Ca^2+^ stores are another source of Ca^2+^ transients^15^. These local sources of Ca^2+^ rise are generally driven by incoming inputs and do not always result in action potential outputs. Especially in awake behaving animals, neurons are spontaneously active in most of brain areas, although they are not related to behaviors. These intrinsically occurring background signals will cause Ca^2+^ rise at a number of synapses and dendritic branches eventually causing aggregation of gene expression at the cell body. Particularly when Cal-Light is applied *in vivo* using viral infection which requires several weeks for sufficient expression of Cal-Light constructs, non-specific signals may accumulate over time, resulting in decreased specificity.

To resolve these weaknesses, we developed a soma-targeted version of Cal-Light (ST-Cal-Light). By concentrating the expression of Cal-Light constructs in the cell body, the system mainly converts Ca^2+^ signals caused from somatic action potentials into gene expression by reducing the portion of other Ca^2+^ sources-dependent gene expression. Thus, cell labeling becomes more dependent on action potential numbers at the soma. Furthermore, because a high amount of Cal-Light expression is condensed at the cell body, its responsiveness to light and Ca^2+^ becomes higher, decreasing the labeling time. The reduction in time thus increases the specificity of successful labeling making the Cal-Light technique applicable to a much broader spectrum of behaviors.

The previous Cal-Light also requires the injection of three viruses. Because virus infection rate is not 100%, a mixture of three viruses causes expression variation in cell to cell. To reduce this limitation, here we generated the ST-Cal-Light KI mouse line. Furthermore, because the KI mouse is designed to be conditional, breeding with other genetically defined cell-type specific mice enable targeting of functionally active neurons in a designated cell type.

## Results

### Development and characterization of ST-Cal-Light *in vitro*

We used two different soma-targeting peptides to facilitate the localization of Cal-Light to the cell body membrane. The first was a 150 amino acids fragment at the amino-terminus of kainate receptor 2 (KA2)^16^ and the other was a 65 amino acids fragment at the carboxyl-terminus of Kv2.1^16,17^ To test whether these two motifs restricts Cal-Light expression to the cell body, we inserted the soma-targeting fragment between the cytosolic side of transmembrane (TM) domain and calmodulin (CaM) of the main Cal-Light construct (Myc-TM-*KA2 or Kv2.1 motif*-CaM-TEV-N-AsLOV2-TEVseq-tTA) (Fig. 1a). Another Cal-Light component, M13-TEV-C, is allowed to freely diffuse in the cytosol without having any localization motif, so that it can interact with the other partner without spatial restriction.

**Fig. 1.**
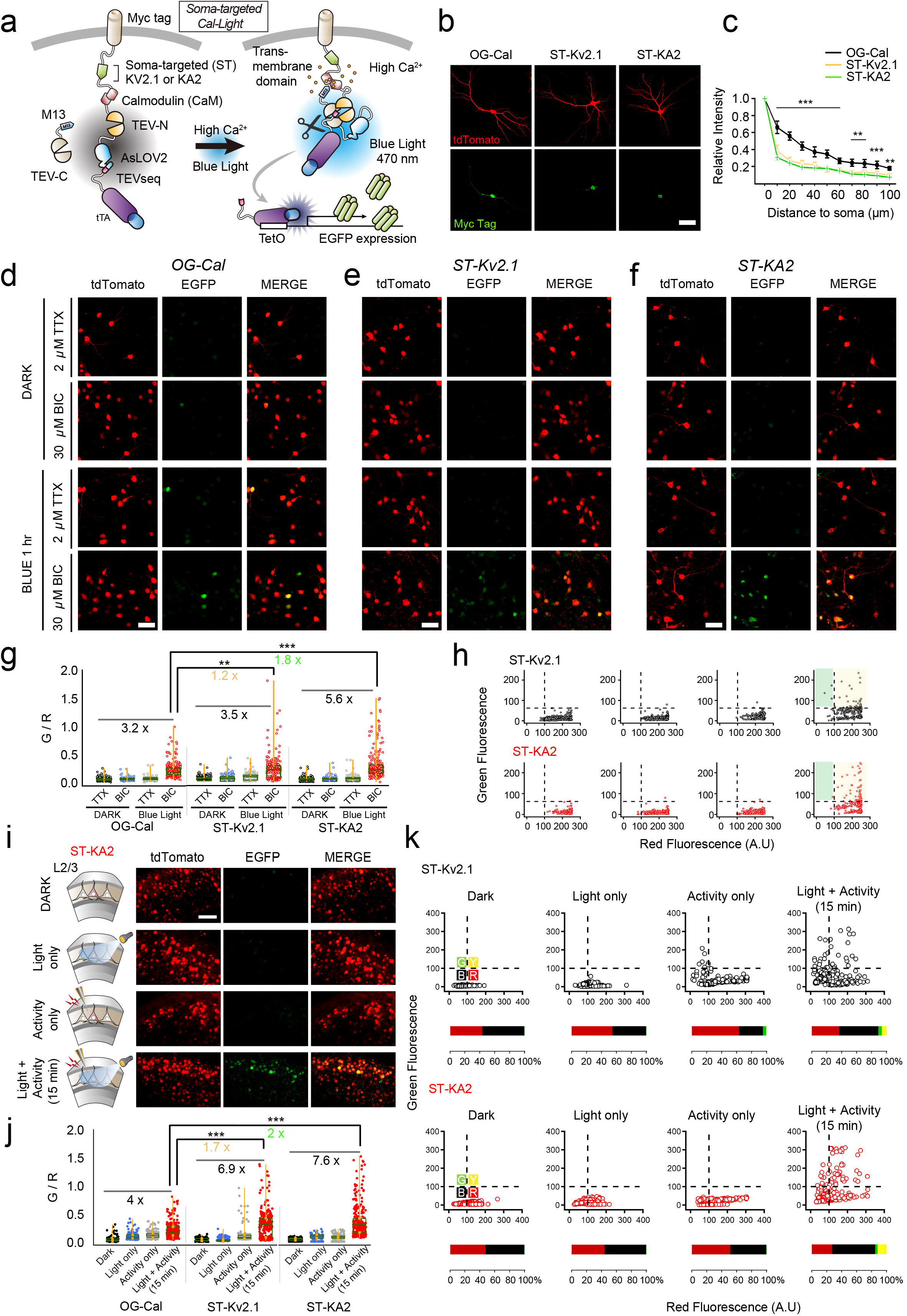
Development and verification of soma-targeted Cal-Light. **a**, Graphical illustration of soma targeted Cal-Light system. Elevation of Ca^2+^ concentration in the cytosol causes M13 and CaM protein interaction which causes binding of c- and n-terminus of tobacco etch virus (TEV) protease (TEV-C and TEV-N). When TEV-C and –N fragment bind, they regain proteolytic functions; however, TEV recognition sequence (TEVseq) is buried at the inside of AsLOV2 Jα-helix, so TEV protease access to the TEVseq is prohibited. Blue light triggers structural changes of AsLOV2, rendering the TEVseq exposed to the cytosol. Then, TEV protease cleaves out tTA, and gene expression begins. **b**, Confocal images of the cell transfection marker, tdTomato (red), and antibody staining of myc epitope (green). Scale bar indicates 50 μm. **c**, The degree of expression in soma and dendrites. Green to red signal (G/R) ratio was normalized to the cell body and the ratio was measured from the cell body up to 100 μm in dendrites. **d-f**, Representative images of EGFP reporter gene expression in various conditions. Original Cal-Light (**d**), ST-KV2.1 (**e**) and ST-KA2 constructs (**f**) were transfected in hippocampal culture neurons and neural activity was controlled by 2 μM TTX or 30 μM bicuculline (Bic). Scale bar indicates 50 μm. **g**, Comparison of gene expression. G/R ratios from individual cells are plotted. Intermittent flash of blue light (1 sec ON/9 sec OFF) was illuminated for 1 hour. (OG-Cal, Dark TTX: 0.074 ± 0.002, n = 309; Dark BIC: 0.081 ± 0.003, n = 312; Blue TTX: 0.084 ± 0.003, n = 308; Blue BIC: 0.239 ± 0.01, n = 291; ST-Kv2.1, Dark TTX: 0.072 ± 0.002, n = 289; Dark BIC: 0.08 ± 0.003, n = 296; Blue TTX: 0.085 ± 0.004, n = 291; Blue BIC: 0.242 ± 0.012, n = 277; ST-KA2, Dark TTX: 0.06 ± 0.002, n = 280; Dark BIC: 0.058 ± 0.003, n = 262; Blue TTX: 0.065 ± 0.003, n = 270; Blue BIC: 0.299 ± 0.016, n = 250; 5 independent cultures). G/R was analyzed with one-way-ANOVA (p < 0.001). Asterisks (** p < 0.01 and *** p < 0.001) indicate Bonferroni post-hoc significance. **h**, Scatter plot of G/R. Open circles indicate individual neurons. Yellow colored area indicates neuronal population with both green and red signals. Green area indicates cells with a green fluorescence alone. **i**, Schematic of experimental conditions. Representative images from each condition (Dark, Light only, Activity only, Light + Activity) are shown. tdTomato is a transfection marker and EGFP is a reporter. Scale bars, 80 μm. **j**, Summary graph of the G/R ratio from cells transfected with OG-Cal, ST-Kv2.1 and ST-KA2, respectively. G/R values from individual neurons and a summary box plot chart are superimposed. The magnitude was robustly enhanced when both light and activity were present. **k**, Green and red fluorescence values from individual neurons were plotted. Individual neurons were divided into four groups (green, yellow, red, and black) by the level of red and green fluorescence. Horizontal bar graph represents the percentage of red, black, green, and yellow groups. For all graphs, *,**, and *** indicate p < 0.05, p < 0.001, and p < 0.005, respectively. Box plots show the median, 25th and 75th percentiles and whiskers show min to max. Error bars indicate s.e.m.

To confirm somatic localization, we performed antibody staining of myc tag, an epitope added at the outer membrane side of the TM domain. We first examined the localization of the original Cal-Light (OG-Cal), which was found to be expressed in both cell body and dendritic branches (Fig. 1b). Addition of the soma-targeting Kv2.1 or KA2 motif, named ST-Kv2.1 and ST-KA2 respectively, caused rapid signal reduction in dendrites, indicating preferred localization at the cell body (Fig. 1b,c). Cell-fill fluorescence (tdTomato) was broadly distributed throughout all processes of neurons. To test whether gene expression is dependent on Ca^2+^ and light, we transfected either OG-Cal, ST-Kv2.1 or ST-KA2 with a TetO-EGFP reporter into hippocampal culture neurons. Na^+^ channel blocker, tetrodotoxin (TTX), was applied to create a no activity condition, whereas GABAA receptor antagonist, bicuculline, was applied to increase overall neuronal activity. In a dark condition, increasing neuronal activity by bicuculline did not increase EGFP reporter gene expression, and TTX also did not further reduce gene expression level (Fig. 1d-g). These results confirmed that neuronal activity alone was not sufficient to induce gene expression if light is absent, but both blue light and bicuculline induced gene expression robustly (Fig. 1d-g).

All three Cal-Light constructs showed robust Ca^2+^- and light-dependent reporter gene expression, but higher SNR was observed in the case of ST-Kv2.1 and ST-KA2 (1.2 fold higher for ST-Kv2.1 compared to OG-Cal, p <0.01; 1.8 fold higher for ST-KA2 compared to OG-Cal, p < 0.005) (Fig. 1g). To examine the pattern of gene expression from individual cells, a scatter plot of red and green fluorescence was analyzed (Fig. 1h). Overall tendency of positive correlation between green and red fluorescence was observed, but exceptionally strong green signals were found in a few neurons with ST-Kv2.1 infection despite weak red fluorescence. This mismatched reporter gene expression was very low in ST-KA2 transfected cells. Thus, ST-KA2 resulted in the highest SNR.

To verify whether similar results are also obtained in brain slices, we infected adeno-associated viruses (AAV) expressing either ST-Kv2.1 or ST-KA2 into organotypic cortical slice cultures. A stimulation pipette was localized at layer 2/3, and a train of action potentials (10 pulses at 20 Hz per minute) was delivered for 15 minutes. Similar to experiments in dissociated cultures, robust gene expression was induced when both electrical stimulation and blue light were present simultaneously, but not by either individually. The ratio of green to red fluorescence was significantly higher in ST-Kv2.1 and ST-KA2 compared to the OG-Cal (1.7 fold higher for ST-Kv2.1 compared to OG-Cal, p < 0.005; 2 fold higher for ST-KA2 compared to OG-Cal, p < 0.005) (Fig. 1j). The overall fluorescence plot revealed that ST-KA2 resulted in almost no background EGFP reporter gene expression, but a small amount of light-independent expression (resulting from activity only) was found in neurons expressing ST-Kv2.1 (Fig. 1k). Because of lower background signals and high induction capability, we decided to further test ST-KA2 constructs in behaving animals.

#### Labeling and manipulation of active neurons underlying lever pressing behavior

To determine whether ST-KA2 can selectively label behaviorally specific neuronal populations, we trained mice for a lever pressing task as described previously^8^ (see Methods). AAV-ST-KA2 was bilaterally injected into layer 2/3 of the primary motor cortex (M1) (Fig. 2a). Briefly put, water-restricted mice underwent continuous reinforcement (CRF) during which the mice learn lever pressing is associated with water rewards (Fig. 2b). The next sessions are fixed ratio (FR) training with gradual increase in the number of lever presses required to receive a reward. For labeling, we shined blue light (5 sec) whenever the mouse pressed a lever, but then the blue light was prohibited for 25 seconds, even if the mouse presses a lever continuously (5 sec ON/25 sec OFF cycle) (Fig. 2c). To obtain the maximum level of labeling, labeling protocol started from CRF session 15 (CRF15) to FR session 12 (FR12) (Full label). To compare with a weak labeling protocol, we also tested a mild labeling process in which blue light was illuminated only during FR 8-12 (Fig. 2b). The total blue light exposure time was about 550 seconds for full labeling and 350 seconds for mild labeling (Fig. 2g). These protocols, especially for full labeling, were performed in order to match the same conditions in which the previously developed Cal-Light technique was tested^8^. In both cases, gene expression was strongly induced, but the full labeling condition showed more robust gene expression (Fig. 2e, f).

**Fig. 2.**
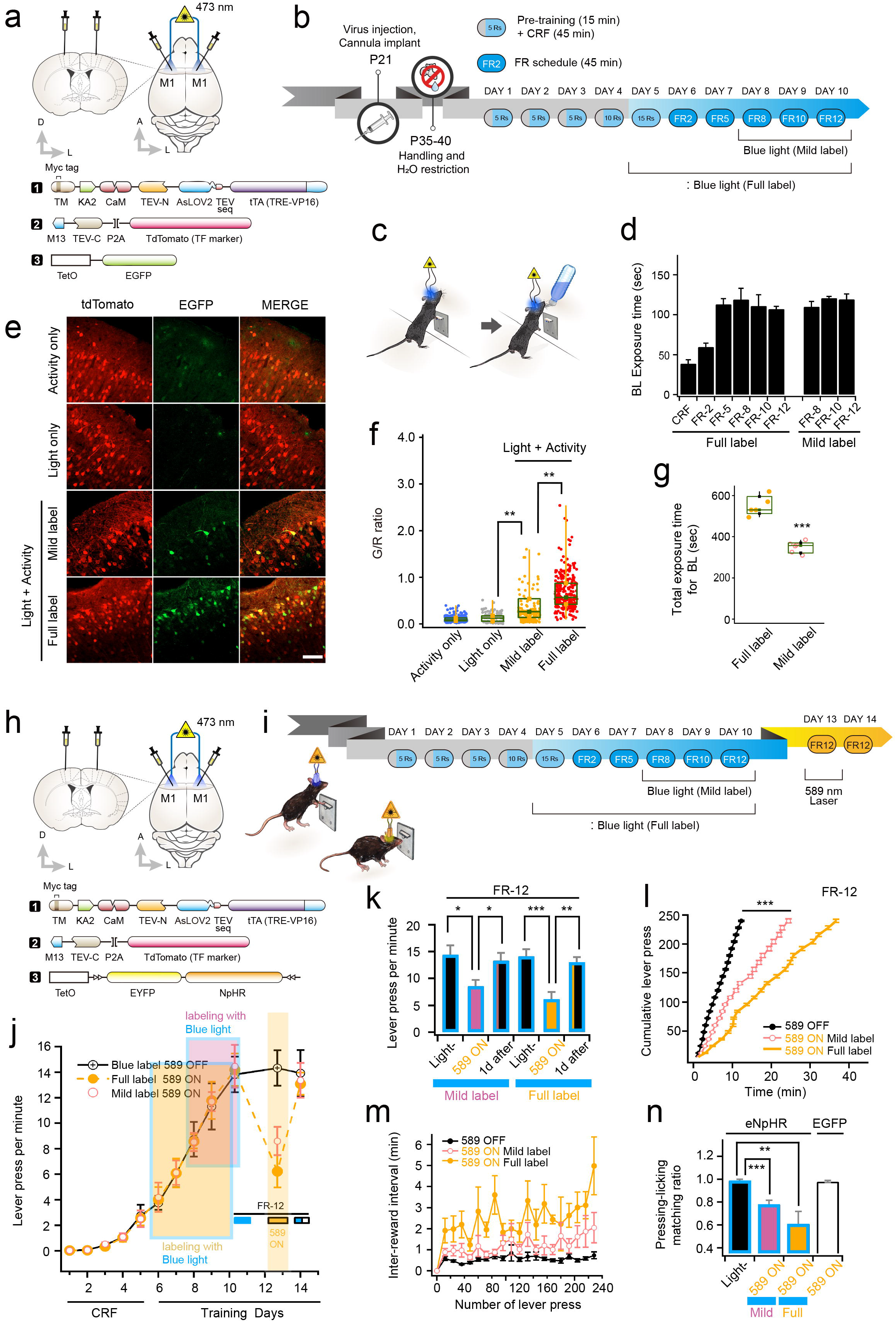
Labeling and control of active neurons engaged with lever pressing behavior. **a**, ST-KA2 viruses with AAV-TetO-EGFP were bilaterally injected into the layer 2/3 of the primary motor cortex. **b**, Schematic mouse training schedule and labeling procedures with blue light. **c**, Fiber optics for blue light illumination were implanted in both sides of the primary motor cortex. To label neurons related to lever pressing,, the laser is turned on for 5 sec whenever the mouse presses the lever followed by a 25 sec time out period in which this laser will not turn on regardless of lever pressing behavior. **d**, Blue light exposure time per training day was measured. **e**, Confocal images showing EGFP expression at each condition. tdTomato signal indicates the efficiency of viral injection. Higher green fluorescence was observed as blue light exposure time increases. Scale bars, 100 μm. **f**, Individual G/R ratios with the box plot chart superimposed. **g**, A box plot chart for total blue light exposure time at each condition (Full, mild label). **h**, Schematic drawing of virus injection and fiber optic implantation (top) and injected three viral constructs (bottom). **i**, Mouse training, labeling by blue light, and halorhodopsin inhibition procedures. Active neurons during lever pressing were labeled by blue light and their activity was inhibited by 589 light. **j**, Total number of lever presses increased over training days, indicating learning. Periods of blue light labeling were indicated by shaded boxes with different colors (5 mice for full labeling, 6 mice for mild labeling). The number of lever presses was significantly reduced by 589 nm laser but fully recovered the following day in the absence of 589 nm light. **k**, Average of lever press number per minute under each condition. **l**, Cumulative lever press number plotted against time. **m**, Inter-reward interval was prolonged when yellow light was turned on. **n**, Summary graph of lever pressing-licking matching ratio. For all graphs, *,**, and *** indicate p < 0.05, p < 0.001, and p < 0.005, respectively. Box-and-whisker plot shows the median, 25th and 75th percentiles and whiskers show min to max. Error bars indicate s.e.m.

To determine whether altering activity of the labeled neurons is sufficient to perturb the learned lever pressing behavior, we injected ST-KA2 viruses together with a halorhodopsin (NpHR) reporter (Fig. 2h). Two days after the labeling procedure, we delivered 589 nm light to suppress activity of the labeled neurons by activating NpHR (Fig. 2i). The number of lever press was counted as an indication of learned behavior (Fig. 2j). Once mice finished FR12 training, 589 nm light (2 sec ON/1 sec OFF) significantly reduced the lever pressing number regardless of full or mild labeling (Fig. 2j). These results indicate that labeling with a shorter period of blue light illumination (~ a few minutes) was enough to prevent learned behavior. Normal lever pressing behavior was fully recovered on the following day, suggesting that the perturbed behavior was not due to the tissue damage or long-term circuit changes by the yellow laser (Fig. 2j, k). Consistent with the reduced number of lever presses, a longer time was needed to reach 250 lever presses and inter-reward intervals were prolonged in the presence of 589 nm light (Fig. 2l, m). Even if mice pressed the lever, the significant portion of lever pressing events was not coupled to water rewards, reflecting the erasure of memory linking lever pressing to rewards (Fig. 2n). When an EGFP reporter was used as a negative control the 589 nm light did not impair lever pressing behavior, verifying that behavioral suppression was indeed caused by NpHR activation, not by the yellow light itself (Fig. 2n).

#### Application of the ST-Cal-Light in short-lived behaviors

Lowering background noise signals while maintaining high inducibility of gene expression may allow for cell labeling during a short period of time. This is critical for *in vivo* application because many behaviors, such as social interaction, are transient and not recurrent. To test if ST-KA2 can be applied for such short-lived behaviors, we assessed several types of behaviors. We injected a mixture of viruses expressing ST-KA2, M13-TEV-C, and TetO-NpHR-EYFP into the CA1 region of the dorsal hippocampus (Fig. 3a). The neuronal population engaged with contextual fear memory was labeled by shining blue light concomitantly with a foot shock. This paring was repeated 3 times (1 min interval), so the total blue light exposure was 15 seconds (Fig. 3a). Two days after fear acquisition, conditioned mice displayed freezing responses during the retrieval phase as anticipated; however, 589 nm light robustly reduced the percentage of freezing (Fig. 3b). We confirmed robust NpHR gene expression in a broad area of hippocampal CA1 (Fig. 3c-e). Thus, ST-KA2 efficiently labeled cell populations encoding conditioned fear memory through minimal bouts of light flashes, and their behavioral causality was verified.

**Fig. 3.**
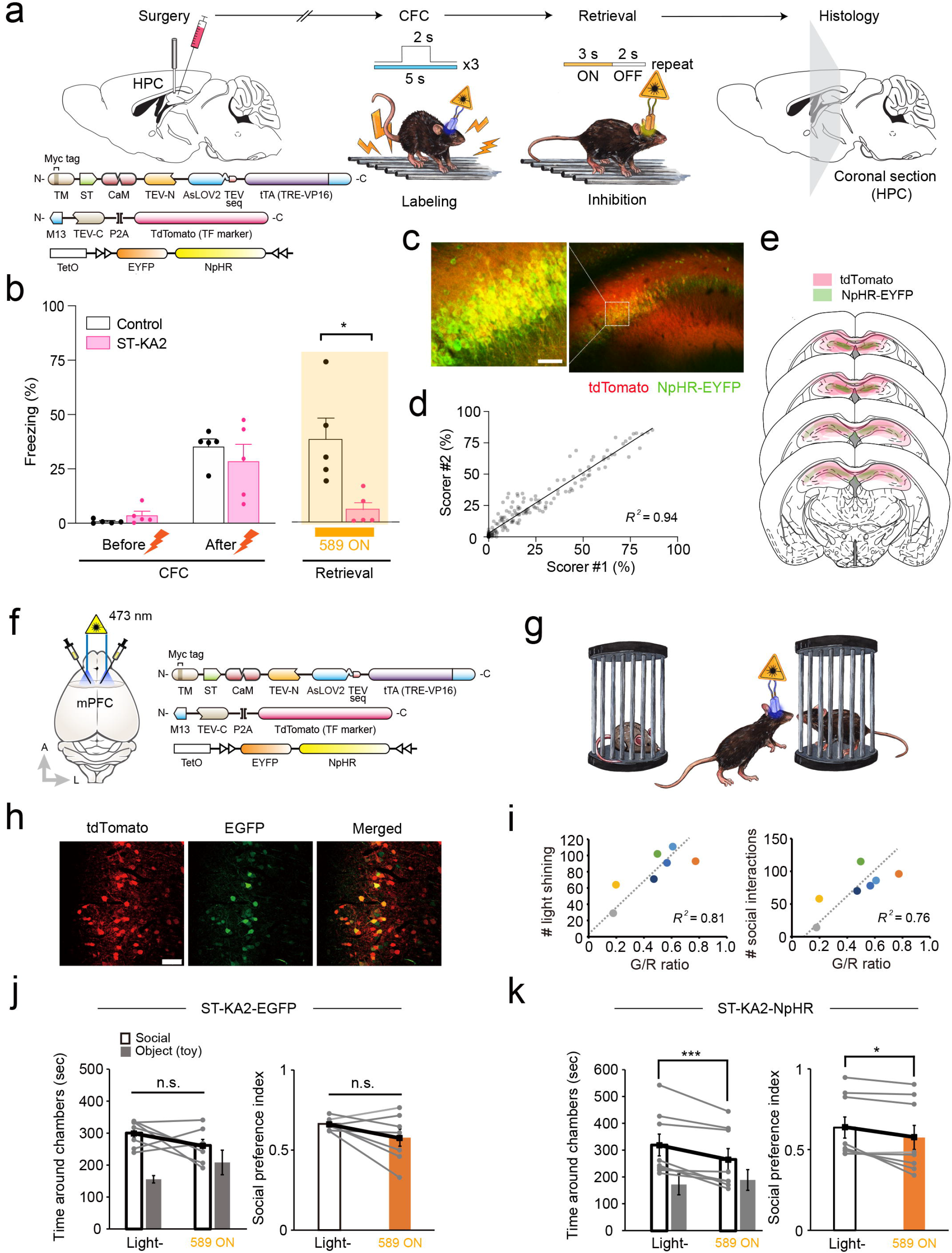
Controlling context-dependent fear conditioning and social interaction behavior. **a**, Schematic illustration of virus injection and fear conditioning experiments. A mixture of viruses expressing ST-KA2, M13-TEV-C, and TetO-NpHR-EYFP into dorsal hippocampus CA1 area. Fiber optics were implanted above the virus injected area. A short pulses of blue light (5 sec x 3 times) were delivered for labeling active neurons and yellow light was shined during the probe test. Reporter gene expression was confirmed by taking confocal images. **b**, The percentage of freezing behavior was compared before and after conditioning, and with or without 589 nm light during retrieval period. (CFC; *N*_control_=5, *N*_ST-KA2_=5; two-way repeated measures ANOVA, AAV x Shock, *F*_1,8_=1.474,*p* 0.259, Retrieval; *N*_control_=5, *N*_ST-KA2_=5; two-tailed student’s t-test, *t*_8_=3.118,*p*=0.014). *\*p* < 0.05. Graphs expressed as mean ± SEM. **c**, Representative image of NpHR-EYFP expression. Scale bars, 50 μm. **d**, Freezing score was analyzed by two independent people in a blind manner. The freezing percentage was scored every 10 s, and crossed-checked with correlation analysis. **e**, The extent of virus injection and NpHR-EYFP expression is plotted across several coronal sections of brain. **f**, Virus injection and fiber optic implantation scheme. Viruses were injected into layer 5/6 of the mPFC. **g**, Cartoon for social interaction experiments. Whenever mouse entered a social zone, a blue laser connected to the fiber optics was switched on. **h**, Sample images of active neuron labeling in the mPFC by ST-Cal-Light. Scale bars, 100 μm. **i**, Graphical demonstration of positive correlation between the number of blue light illumination/social interactions and G/R ratio. **j**, When active neurons were labeled with EGFP reporter, 589 nm light did not decrease social interaction (Light -: 300 ± 14.7 sec, 589 nm ON: 261 ± 18.9, n = 7, p > 0.05) **k**, NpHR reporter gene was expressed during labeling process, social interaction behavior was significantly inhibited during the probe test (Light -: 318 ± 40.3 sec, 589 nm ON: 264 ± 38.3, n = 8, p < 0.05).

We further tested whether ST-KA2 can also target neurons involved in behaviors based on high cognitive functions. The medial prefrontal cortex (mPFC) has been well characterized for mediating social cognition^18–22^, but functionally involved neural populations have not been analyzed at an individual cell resolution. To test whether ST-KA2 can selectively label an mPFC neuronal population engaged with social interaction, a mixture of AAVs expressing ST-KA2, M13-TEV-C, and TetO-EGFP were bilaterally injected into the layer 5/6 of the mPFC (Fig. 3f). For reporter gene induction, a blue light was programmed to be switched on for 5 seconds whenever the test mouse entered a social zone (3 cm from the social box) to interact with another mouse (Fig. 3g). Because of mouse to mouse variability in interaction time, the amount of blue light exposure was different in each animal. *Post-hoc* confocal imaging revealed that EGFP reporter gene expression was present in a subset of neurons (Fig. 3h). Additionally, EGFP expression was positively correlated with the amount of blue light exposure (Fig. 3i). These results confirmed that the magnitude of gene expression mediated by ST-Cal-Light is dosedependent. To confirm the behavioral causality of the labeled neurons, we shined a 589 nm light in animals injected with an NpHR reporter vector. Yellow light reduced the degree of social interaction as indicated by a reduced social preference index (Fig. 3k). The control group administrated with the same yellow light and the same viruses with an EGFP reporter did not cause any changes in social interaction behavior (Fig. 3j), suggesting that reduced social interaction was not due to the yellow light itself. Thus, the ST-Cal-Light could specifically visualize and confirm the behavioral causality of neurons governing social interaction.

#### Application of the ST-Cal-Light in brain diseases

We next tested whether ST-Cal-Light can also be used to dissect neuronal population underlying specific brain disorders. It is generally believed that epileptic seizure is caused by unobstructed electrical discharge in the hippocampus^23^. However, alleviating behavioral symptoms by directly targeting only engaged neurons has not been directly tested. A mixture of viruses (AAV-ST-KA2, AAV-M13-TEV-C-P2A-TdTomato, and TetO-EGFP) were injected in hippocampal dentate gyrus (DG) and CA1 areas (Fig. 4a). A seizure was induced by administrating kainic acid (KA) intraperitoneally (20 mg/kg) (Fig 4b). Shortly after KA injection (10 min), blue light (3 sec ON/2 sec OFF, 30 min) was illuminated in order to label active neuronal populations linked to epileptic seizure. In a control group, the same amount of KA was administered, but blue light was not delivered to determine whether labeling is light-dependent or not. As predicted, robust gene expression was only present in the group which received blue light illumination (Fig 4c). If labeled neurons are directly coupled to behavioral abnormalities, we may be able to ameliorate seizure symptoms by selectively suppressing activity of labeled neurons. For this experiment, an AAV expressing the TetO-NpHR reporter was injected together with AAV-ST-KA2, and AAV-M13-TEV-C-P2A-TdTomato (Fig. 4d). After ST-Cal-Light proteins were fully expressed, the seizure-specific neurons were labeled during the first KA injection. A second KA injection was tested either with or without shining 589 nm light (Fig. 4e). The severity of seizure phenotypes was analyzed by division into multiple stages including immobilization, head nodding, continuous myoclonic jerk, and clonic-tonic convulsions (see Methods). Symptoms were exacerbated over time after systematic KA injection, but delivery of 589 nm light significantly suppressed the progression of seizure symptoms (Fig. 4f-h). Inhibiting hippocampal neurons involved in context-dependent fear conditioning did not reduce seizures following KA injection (Fig. 4h). These results suggest that seizure symptoms are mediated by specific cell population, but not by nonspecific hippocampal neurons. Post-hoc staining showed that granule cells in the DG, mossy cells in the hilus, and CA1 and CA3 hippocampal neurons were labeled (Fig. 4i). Because the ST-Cal-Light is designed to express higher NpHR expression in more active neurons, it is presumed that highly active neurons such as superhub neurons^24,25^ were preferentially inhibited in our experiments. Thus, the ST-Cal-Light would be useful for dissecting neural circuits engaged in brain disorders at the individual cell or subpopulation level.

**Fig. 4.**
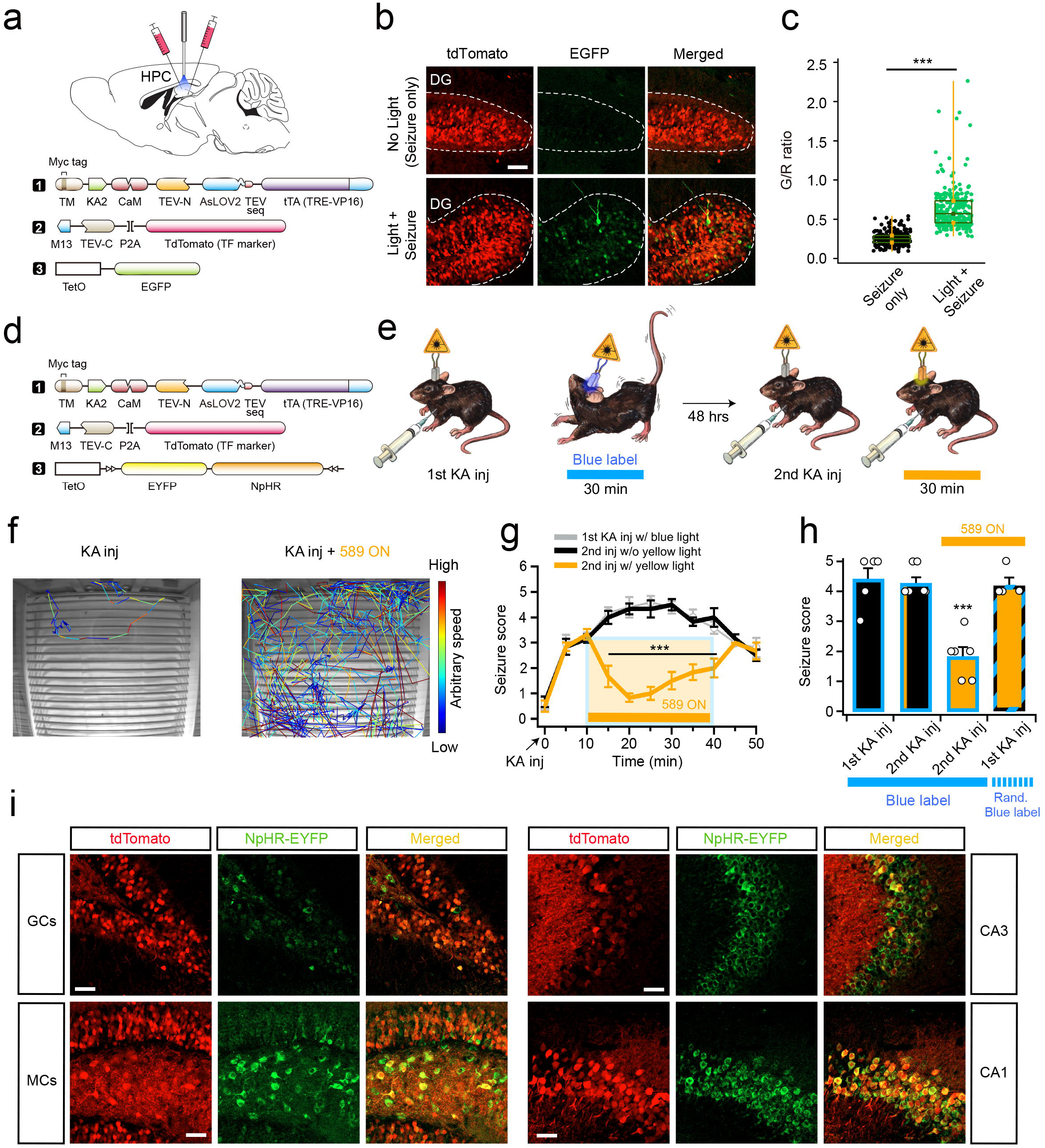
Amelioration of epileptic seizure by ST-Cal-Light. **a**, Schematic drawing of virus injection and fiber optic implantation. ST-Cal-Light viruses with TetO-EGFP reporter were injected into both hippocampal CA1 and CA3 areas bilaterally. **b**, Representative images of tdTomato (transfection marker) and EGFP reporter gene expression. When seizure was induced by KA administration, little gene expression was created in the absence of light, demonstrating cell labeling was dependent on blue light. **c**, A box-and-whisker plot of G/R ratio. **d**, For inhibition experiments, a TetO-NpHR reporter was used. The same ST-Cal-Light viruses were injected as shown in **a**. **e**, A cartoon demonstrating experiment procedures. Seizure was induced by KA injection and blue light was illuminated for labeling. Two days later, KA was injected again for the second seizure induction and compared the severity of seizure with or without 589 nm light. **f**, Sample movement traces after KA injection with or without yellow light. Movement during the same period of time was plotted. Different colors represent animal movement speed. ezTrack analysis was used for tracing (from Denise Cai lab)^33^. **g**, Time lapse changes of seizure score after KA administration. **h**, Average seizure scores at various conditions. **i**, Hippocampal granule cells (GCs), mossy cells (MCs), CA1 and CA3 neurons were labeled by ST-Cal-Light, indicating increased neuronal activity in broad hippocampal areas.

#### Generation of conditional ST-Cal-Light knock-in mice

Identification of behaviorally relevant neurons within a selective cell type will improve understanding of the specificity of their roles underlying behaviors. To achieve this goal, we generated a conditional ST-Cal-Light KI mouse (Fig. 5). The ST-KA2 gene was inserted into the GtROSA26 locus of the chromosome (Fig. 5b). A LoxP flanked neo cassette was placed in the upstream of target gene, so that cre recombinase can remove the neo cassette, resulting in ST-KA2 expression. To confirm that floxed ST-Cal-Light causes reporter gene expression in a cre-dependent manner, we compared EGFP reporter expression. When AAV expressing cre was not injected to the floxed ST-Cal-Light KI mice, repeated shining of blue lights (3 sec ON/2 sec OFF, 30 min) did not cause EGFP expression (Fig. 5c). In contrast, robust EGFP expression was induced in mice injected with cre-expressing virus, verifying the KI system works in a cre-dependent manner (Fig. 5c). To make ST-Cal-Light system work in all neurons transfected with M13-TEV-C, we constructed a new viral plasmid expressing both M13-TEV-C and Cre recombinase (Fig. 5d). Introducing this construct with an EGFP reporter showed clear EGFP expression (Fig. 5d). When generating the conditional floxed ST-Cal-Light KI mouse, a myc epitope was added at the outside of TM domain, so antibody staining against myc showed signals in the outer cell membrane (Fig. 5d).

**Fig. 5.**
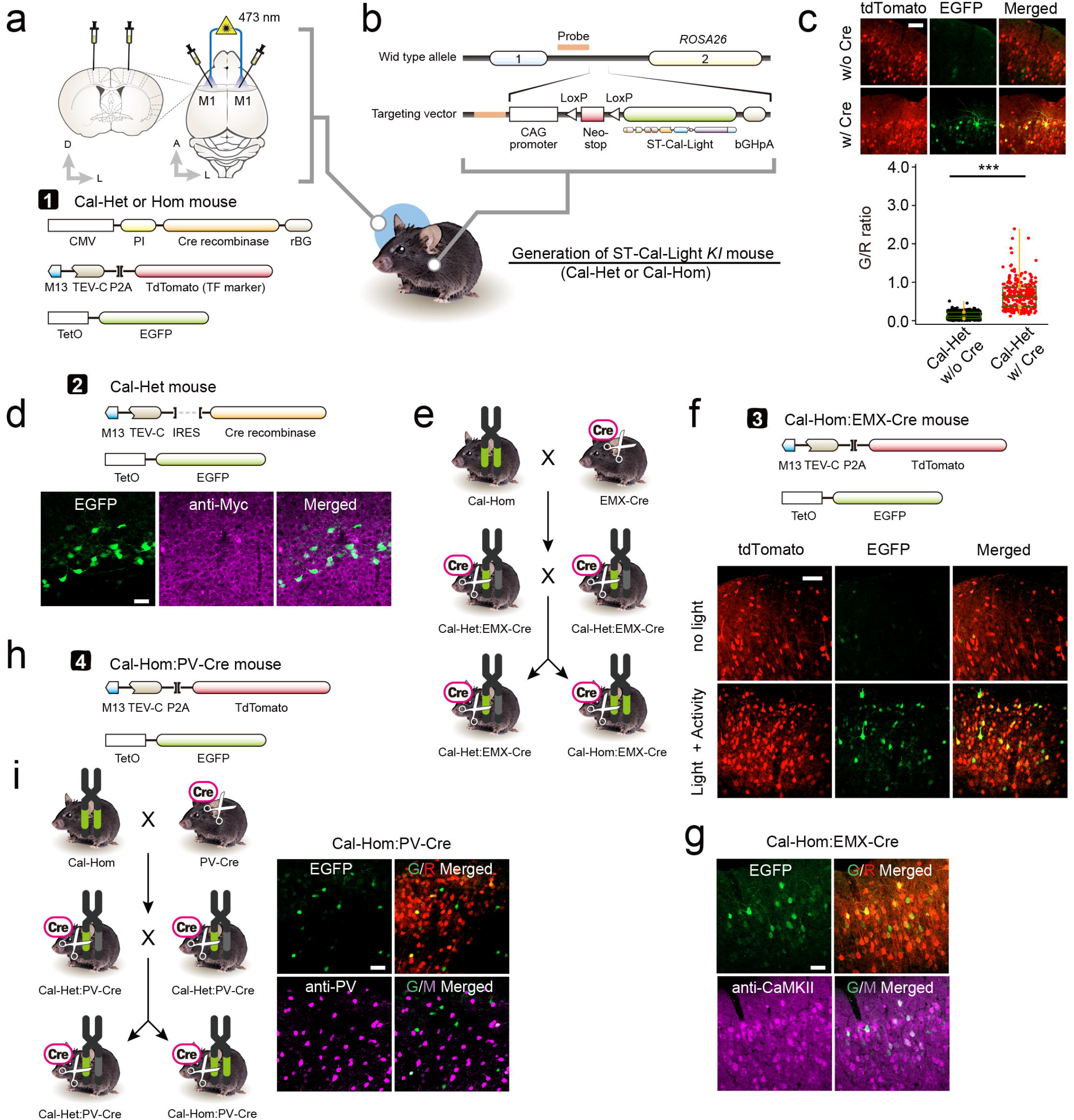
Generation of conditional ST-Cal-Light knock-in mice. **a**, Schematic illustration of virus injection and fiber optic implantation. **b**, Plasmid design for generating conditional ST-Cal-Light KI mouse. **c**, Representative images of tdTomato (transfection marker) and EGFP (reporter) and G/R ratio distributions with or without introducing Cre recombinase. Scale bar, 100 μm. **d**, Injected viruses (top) and images of EGFP and anti-myc staining. Myc epitope is expressed in a cre-dependent manner and localized at the somatic membranes. EGFP signals indicate active neurons. Scale bars, 50 μm. **e**, Cartoon of breeding scheme. **f**, A mixture of viruses were injected into the primary motor cortex of Cal-Hom:EMX-Cre mouse (top). Blue light-dependent gene expression in neocortical excitatory neurons under Emx1 promoter (bottom). Scale bars, 50 μm. **g**, Excitatory neuron labeling was confirmed by CaMKII antibody staining. Scale bar, 50 μm. **h**, Virus injection scheme in Cal-Hom:PV-Cre mice. **i**, Schematic flow of generating either Cal-Het:PV-Cre or Cal-Hom: PV-Cre (left). Active PV-positive neurons were labeled and confirmed by PV antibody staining. Scale bars, 50 μm.

To test cell type specific labeling, we crossed the floxed ST-Cal-Light mouse with Emx1-Cre. We injected AAV-M13-TEV-C-P2A-TdTomato and AAV-TetO-EGFP into the layer 2/3 of primary motor cortex of either ST-Cal-Light hetero- or homozygotes (Fig. 5e). tdTomato signal is driven by CAG promotor, so its expression was not restricted in specific cell types, but EGFP expression was induced in excitatory neurons because Cre expression is limited in neocortical excitatory neurons under Emx1 promoter^26^ (Fig. 5f). Antibody staining with CaMKII, an excitatory pyramidal neuron marker, showed colocalization of CaMKII and EGFP signals (Fig. 5g). Similarly, we tested labeling of active interneurons by crossing the floxed ST-Cal-Light mouse with a PV-Cre mouse (Fig. 5h, i). Repetitive blue light (3 sec ON/2 sec OFF, 30 min) induced EGFP reporter gene expression restricted to PV interneurons (Fig. 5i). Thus, conditional ST-Cal-Light mice specify active neurons from a distinct cell type.

### Discussion

Labeling and manipulating active neurons with high temporal resolution is crucial for understanding circuit functions underlying animal behaviors. A dual switch system with “AND” gate logic using Ca^2+^ and light has successfully converted fast Ca^2+^ transients to slow gene expression, allowing for activity control of labeled neurons^7–10^. Despites these advantages, collecting all signals arising from various Ca^2+^ sources causes labeling that may not be directly associated with specific actions. To minimize such noise signals, we intended to condense the Cal-Light system, restricting it to the cell body. Highly concentrated Cal-Light proteins at the cell soma increases the chance of binding two Cal-Light components, which results in efficient tTA release upon blue light. Thus, soma targeted Cal-Light results in higher expression of reporter proteins while simultaneously reducing off target gene expression. In this scheme, the magnitude of gene expression remains high, but background signals are reduced. Furthermore, because somatic action potentials are typically final output signals of neurons, their inhibition has a more prominent effect on behaviors. All these improvements collectively make the labeling process more efficient as well as shortening the duration and repetition of blue light illumination.

In this study, we examined the efficiency of ST-Cal-Light in four different behaviors. The first was a lever pressing behavior, in which animals press a lever repetitively in order to acquire rewards. Tagging active neurons during such repetitive behavior is one good case for using ST-Cal-Light technique. Because the strength of labeling positively correlates with repeated light exposure and the amount of Ca^2+^ release, more active neurons accrue greater gene expression with each behavioral repeat, while less active neurons will not. Labeling efficiency is maximized if repeated behavior has an inter-trial interval because 30-60 seconds are required after blue light illumination for structural restoration of AsLOV2 domain^27^

In case of transient behaviors, capturing involved neurons requires a high induction rate with low background signals. Our ST-Cal-Light is designed to have concentrated constructs at the cell body, so fewer light repeats can yield higher levels of reporter gene expression. We tested context-dependent fear conditioning because it can be achieved by few trials with very strong stimuli. We found that just three rounds of blue light exposure was sufficient to express NpHR in labeled neurons which, when activated, was sufficient to inhibit the fear memory. Presumably, Ca^2+^ levels were robustly enhanced by electric shock, increasing the efficiency of gene expression. Tagging neurons involved in social interaction is also a good candidate for modeling transient behaviors. Time for single social interaction event is brief but repetitive. The frequency of interaction is also variable between animals. Thus, dissecting out only active cells and testing behavioral causality can be performed from individual animals and correlation between labeling efficacy and behavior can be compared. Because social interaction is not simply mediated by neurons in a single brain area but by multiple circuits^28–32^, viral injections in several brain areas will target involved neurons more comprehensively and optogenetic reconstruction of full social behavior may be possible.

Generation of an ST-Cal-Light KI mouse can improve the degree of specificity by cell type when it combined with other specific cre driver lines. Development of ST-Cal-Light mouse also reduces expression errors or variation caused by virus injection. Tagging active neurons only from a specific cell type will give a better understanding about circuit functions. We confirmed that the conditional ST-Cal-Light mice limited gene expression only to specific cell types. Thus, the ST-Cal-Light system allows investigators to dissect neural circuits into individual cells and test direct behavioral causality in space and time.

## Data availability

All plasmids used in this study were deposited and available in Addgene (ID: 171615 ~ 171622). The conditional ST-Cal-Light KI mice are available from the corresponding author upon request.

## Acknowledgements

We thank members of the Kwon laboratory for helpful discussion, and Holly Robinson for manuscript proofreading. This work was supported by the National Institute of Health Grants DP1MH119428 (to H-B.K.), NS036715 (to R.L.H.), MH-092443 (to A.S.), MH-094268 (to A.S.), MH-105660 (to A.S.), and MH-107730 (to A.S.); foundation grants from Stanley (to A.S.), and RUSK/S-R (to A.S.). This work was also supported by National Research Foundation of Korea (NRF) grants (to D.L.); the Korea Government Ministry of Science and Information and Communications Technology (ICT) (NRF-2018M3C7A1024597) (to D.L.).

## Author contributions

H-B.K., J.H.H. and D.L. conceived and designed the study. J.H.H. performed experiments and data analysis related to lever pressing training, seizure induction, and conditional KI mouse verification. J.H.H. did data analysis from slice culture experiments. K.N. performed social interaction experiments and related data analysis. H.N. and A.S. performed experiments and data analysis related to context-dependent fear conditioning. N.M. performed dissociated cell culture experiments. P.H. helped surgeries and behavior experiments related to lever pressing. S.K. performed experiments using organotypic slice culture. M.C., S.L., D.L. performed DNA plasmid construction and cloning. B.L. and R.H. performed the initial verification of myc antibody staining. A.M. helped slice culture data analysis. C.K. performed behavioral analysis using the ezTrack and DeepLabCut method. H-B.K. wrote the first draft and all authors edited the manuscript.

## Corresponding author

Correspondence to Hyung-Bae Kwon.

## Ethics declarations

Competing interests

The authors declare no competing financial interests.

## Methods

### Construction of plasmids

Entire DNA sequences of constructs used in this study are described in the Supplementary information. Concisely, to generate pAAV::CMV-FLEX-TM-KA2-CaM-TEV-N-AsLOV2-TEVseq-tTA, pCMV::TM-KA2-CaM-NES-TEV-N-AsLOV2-TEVseq-tTA, and pCMV::TM-KV2.1-CaM-NES-TEV-N-AsLOV2-TEVseq-tTA, we acquired DNA sequences for CaM, TEV-N, AsLOV2, and tTA from pAAV::TM-CaM-NES-TEV-N-AsLOV2-TEVseq-tTA (Addgene #92392) by the conventional PCR reactions. Amplified PCR products and digested pAAV or pCMV empty backbone were cloned in the overlap cloning technique using overlap cloner (ELPIS-BIOTECH, CAT #EBK-1012). Similarly, pAAV::M13-TEV-C-IRES-SP6-Cre was constructed by inserting amplified M13 and TEV-C, and synthesized IRES-SP6-Cre into the pAAV empty vector. To build pCMV::TM-KA2-CaM-NES-TEV-N-AsLOV2-TEVseq-tTA and pCMV::TM-Kv2.1-CaM-NES-TEV-N-AsLOV2-TEVseq-tTA, both KA2 and Kv2.1 were synthesized in Macrogen and sequentially digested by XhoI, and each fragment was inserted between TM and CaM domain of pCMV::TM-CaM-NES-TEV-N-AsLOV2-TEVseq-tTA. eNpHR-EYFP which was amplified from Addgene #20949 and hChR2 (H134R) from Addgene #20297 using the conventional PCR. Amplified fragments were inserted into AAV backbone vector respectively to clone pAAV::TRE-NpHR-EYFP and pAAV::TRE-ChR2-YFP. All restriction enzymes and reagents for cloning were purchased from New England BioLabs and ELPIS-BIOTECH (Republic of Korea). Cloned plasmid vectors were carefully confirmed by the DNA sequencing analysis (Macrogen).

### Primary neuron culture and DNA transfection

Primary dissociated neuron cultures were performed as described in previous literature (Lee et al., 2012). To briefly describe the process, CD IGS rat hippocampus (embryonic day 18-19) were quickly dissected and digested in 0.25% trypsin-EDTA (Invitrogen) at 37°C for 8-10 minutes. Trypsin-EDTA was then removed and the hippocampal brain tissue was gently triturated ~10-15 times using a 100-1,000uL pipette tip. 12-mm PDL-coated coverslips (Neuvitro) were placed on 24-well plates. Dissociated cells were counted and plated with a 10^5^ cells concentration in each well. The medium used to plate neurons consisted of neurobasal medium (Invitrogen) with 1% FBS (Thermo Fisher Scientific), 1% Glutamax supplement (Gibco), and 2% B27 supplement (Gibco). Every 3-4 days, one-half of the media was replaced with freshly prepared medium lacking FBS. On DIV 9, cultures were transfected with DNA constructs using a Lipofection method (Thermo Fisher Scientific, Lipofectamine 3000 Reagent, #L3000008) for sparse DNA transfection. Diluted DNA constructs were mixed into the Lipofectamine reagent at a 1:1 ratio for each membrane specific receptors (pCMV-TM-*KA2 or Kv2.1*-CaM-NES-TEV-N-AsLOV2-TEVseq-tTA and M13-TEV-C-P2A-tdTomato). On DIV 14, all neuronal cultures were fixed and imaged using a confocal microscope (Zeiss LSM880).

### Myc tag staining *in vitro*

Myc (to visualize soma targeted receptor) and tdTomato (for M13-CaM transfection confirmation) staining for neuronal cultures was accomplished as follows: individual cultures were rinsed three times in PBS, pH 7.4; slices were blocked in 10% normal donkey serum (Jackson ImmunoResearch, 017-000-121) and 0.1% Triton-X for 30 minutes and inserted into a shaking incubator set to 23°C at 120-130 RMP; incubated with a mixture of RFP antibody preabsorbed (1:1000 in blocking reagent, Rockland antibodies & assays, 600-401-379) and Anti-Myc tag antibody (1:1000 in blocking reagent, Abcam, #ab32) for 90 minutes in room temperature shaking incubator; rinsed in PBS three times with 5-minute incubation periods each time; incubated in Cy3-AffiniPure Donkey Anti-Rabbit IgG (H+L) (1:500, Jackson ImmunoResearch, 711-165-152) and Alexa Fluor 488 AffiniPure Donkey Anti-Mouse (1:500, Jackson ImmunoResearch, 715-545-150) for 30 minutes at room temperature shaking incubator; rinsed with PBS three times with 5-minute incubation periods; mounted with DAPI Fluoromount-G (Southern BioTech, 0100-01). All cultures were imaged with a confocal microscope (Zeiss LSM880).

### Measuring the brightness of the fluorescence along the somatodendritic axis

The image analysis was performed through ImageJ software. After defining the boundaries of the soma from each neuron, a 20 μm diameter circle near the soma was made where no apparent fluorescence exists. The average fluorescence in this circle was used as background fluorescence. The boundary of the soma was defined by using polygon tool, and the average fluorescence inside of it was measured and subtracted the aforementioned background value. This value was defined as ‘soma fluorescence’. To measure fluorescence intensities along the dendrites, 1μm^2^ rectangles on the somatodendritic axis were made at every 10 μm from the soma and measured up to100 μm. The distance between each rectangle and the soma was not correspond to the minimal linear distance from the soma (since dendrites were curved). To calculate the ratio of fluorescence intensity at each position, the fluorescence intensity at each rectangle was measured and background was subtracted. This value was divided by the soma fluorescence and plotted as a function of distances along the dendrites.

### Preparation of cortical organotypic slice culture and virus infection

Organotypic slice cultures were made from P2-P4 C57BL/6 mice (Charles River Laboratory). The general procedures for organotypic slice cultures were performed as previously described^8^. Coronal sections of cortex (thickness, 400 μm) were made by a tissue chopper (The Mickle Laboratory Engineering Co. Ltd., UK). The age of culture is indicated by an equivalent postnatal (EP) day; postnatal day at slice culture (P) + days *in vitro* (DIV). On EP 5-8, cultured slices were infected by adding a 5 μl of solution containing 1 μl of concentrated virus (titer: ~10^13^-10^14^ GC/ml) and 4 μl of slice culture media (pre-warmed at 37°C) to the top of the cortical layers of brain slice placed on porous (0.4 μm) membrane (Millicell-CM; Millipore) for covering whole slice to maximize the infection rate. After infection, all the 6-well dishes that contain cultured slices were returned to the incubator (37°C) with the aluminium foil covered to prevent the unexpected light exposure. Experiments were performed at EP 20, two weeks after viral infection. ST Cal-Light viral AAV vectors were cloned in the lab and viruses were produced at ViGene Bioscience (Rockville, MD, USA).

### Activity- and light-sensitive neuronal tagging in organotypic slice cultures

Cultured slices were taken out from the incubator for induction experiment and superfused with autoclaved artificial cerebrospinal fluid (ACSF) solution containing the followings (in mM): 124 NaCl, 26 NaHCO_3_, 3.2 KCl, 2.5 CaCl_2_, 1.3 MgCl_2_, 1.25 NaH_2_PO_4_, 10 glucose, saturated with 95% O2 and 5% CO2 gas. Concentric bipolar electrode (12.5 μm inner pole diameter, 125 μm outer pole diameter; FHC Inc.) was positioned at layer 2/3 cortical area. Stimulation pulses (100 μs in duration, stimulus intensity, 10-15 V) were generated by a digital stimulator (Master 8, ÄMPI, Israel) and fed into the stimulation electrode via an isolation unit (DS2A, Digitimer Ltd). A blue laser (MBL-F-473 nm-200 mW, CNI, China) coupled to a FC/APC fiber (400 μm, 1 meter long, CNI) was positioned two centimeters above the surface of the cortical slice. Total power from the tip of the fiber was 5-10 mW. The induction protocol used in this study elicited about 75 spikes. This number was much lower than one used in a previous study (~900 spikes) (Lee et al., 2017). Blue light was illuminated for 15 min (10 sec ON / 50 sec OFF) with or without electrical stimulation (Light + Activity vs. Light only). One train (5 pulses at 10 Hz) of electrical stimulation was delivered per min with 10 sec-long blue light. Sample whole-cell patch-clamp recordings were made to measure the number of spikes that elicited by the weaker induction protocol through a MultiClamp 700B amplifier controlled by Clampex 10.2 via Digidata 1440A data acquisition system (Molecular Devices). The pipette solution was made as follows (in mM): 125 K-gluconate, 5 KCl, 10 Na_2_-phosphocreatine, 4 Mg-ATP, 0.4 Na-GTP, 10 HEPES, 1 EGTA, 3 Na-ascorbate (pH = 7.25 with KOH, 295 mOsm).

### Animals

Experimental subjects were used from 6 to 12 weeks old C57BL6J mice (Jackson Laboratory, Bar Harbor, ME, USA, Stock No: 000664). Similar number and ages of both male and female mice were randomly chosen for experiments. All mice for behaviour tests were individually housed in a 12-h dark-light reverse cycle and experiments were performed during the dark cycle period. Mice born into our colony on a C57BL6J background were maintained in conventional housing with free access to food *ad libitum* when not being tested. Cre mouse lines used in this study were purchased from Jackson Laboratory: EMX1-Cre (Stock No: 005628), PV-Cre (Stock No: 017320). All experimental procedures and protocols were conducted with the approval of the Max Planck Florida Institute for Neuroscience (MPFI) Institutional Animal Care and Use Committee (IACUC), Johns Hopkins University IACUC and National Institutes of Health (NIH) guidelines.

### Stereotaxic surgeries

Viral injections, and fiber implantations were performed as described in a previous study^8^ with the following specifications. For targeting primary motor cortex (M1) area, virus containing solution (pAAV-pCMV-Myc-TM-KA2-CaM-NES-TEV-N-AsLOV2-TEVseq-tTA: pAAV-hSYN-M13-TEV-C-P2A-tdTomato: pAAV-TRE-EGFP = 2: 2: 5 ratio) was injected for the labeling of active population of neurons at: ÄP, +0.25 mm; ML, +1.5 mm from bregma, and DV −0.2 mm from the brain surface or pre-mixed viral solution (pAAV-CMV-Myc-TM-KA2-CaM-NES-TEV-N-AsLOV2-TEVseq-tTA: pÄÄV-hSYN-MI3-TEV-C-P2Ä-tdTomato: pAAV-TRE-eNpHR-EYFP, 2: 2: 5) was injected to the same target site for the manipulation of the labeled subset of neurons. After viral injection, the optic fibers (200 μm core; NA 0.37; Cat# BFL37-2000, Thorlabs) were implanted perpendicularly into the targeted brain region. The tip of fiber was positioned 100-150 μm above the target viral injection site in M1 area. To label and manipulate task-relevant neuronal populations in hippocampal CA1 neurons during contextual fear conditioning (CFC) task, the same 500 nl mixture of AAV viral solution was injected into the dorsal hippocampal CA1 region at the following coordinate (AP, −2.0 mm; ML, ±1.3 mm; DV, −2.05 mm from bregma) and the optic fibers were implanted 300 μm above injection site to deliver the light down to the target brain region. For social cognition experiment, we bilaterally injected either of pAAV-CMV-Myc-TM-KA2-CaM-NES-TEV-N-AsLOV2-TEVseq-tTA: pAAV-hSYN-M13-TEV-C-P2A-tdTomato: pAAV-TRE-EGFP (for labeling) or pAAV-CMV-Myc-TM-KA2-CaM-NES-TEV-N-AsLOV2-TEVseq-tTA: pAAV-hSYN-M13-TEV-C-P2A-tdTomato: pAAV-TRE-eNpHR-EYFP (for manipulation) into the mPFC area (AP, +2.0 mm; ML, ±0.5 mm; DV, −1.6 mm from bregma) and the optic fibers were implanted into the same hole of virus injection at 5-10° angle. The tip of the fiber was placed 500-700 μm above sites of the virus injection using a cannula folder (Thorlabs). For *in vivo* seizure experiments, multiple injections per hemisphere were targeted bilaterally to maximize the number of neurons responsible for the seizure activity (CA1: AP, −1.8 mm; ML, ±1.5 mm from bregma, and DV −1.7 mm; DG: AP, −1.9 mm; ML, ±1.1 mm; DV −1.8 mm from bregma). To label the broader areas and the numbers of neurons during seizure, multiple optic fibers were implanted over hippocampus at (relative to Bregma): AP, −1.8 mm; ML, ±1.3 mm; DV −1.2 mm. To relieve post-surgical pain, an analgesic (buprenorphine, 0.05 mg kg^-1^ of body weight) was injected subcutaneously and mice were returned back to their home cage for recovery.

### Optical labeling of the task-related neurons

Three weeks after viral injection, blue light (473 nm) was delivered to the regions of interest (M1, mPFC or hippocampus) during specific behavior. Blue laser (MBL-FN-473, Changchun New Industries Optoelectronics Technology, Jilin, China) was controlled by MED-PC IV software, which connected with an acquisition board (Med-Associates, St. Albans, VT).

#### 1. Skilled motor learning

Water restriction was started 3-4 days prior to the lever-press training and water was restricted throughout the training periods so the mice receive the rest of the water after subtraction of the amount of water given during training to make total 1 ml of water per day. Training was performed in a standard mouse operant chamber (Med-Associates, St. Albans, VT) placed in sound attenuating cubicle (ENV-022MD, 22 cm × 15 cm × 16 cm). In the continuous reinforcement (CRF) session, a mouse received a water reward provided from a retractable sipper tube extended into the chamber after lever press and they learned that lever pressing was associated with water reward. When mice finished getting fifteen rewards (CRF 15Rs), they were moved to the fixed ratio (FR) schedule, which consists of FR-2 (two presses to get a reward), FR-5, FR-8, FR-10 and FR-12 (lever pressing twelve times to get reward once) for 5 consecutive days. These FR sessions lasted 45 min or until mice received 20 rewards. Labeling was performed by two separate protocols. For full labeling, 473 nm blue light (5 sec ON/25 sec OFF cycle) was illuminated from CRF 15Rs to FR-12. For mild labeling, blue light with the same cycle was delivered from FR-8 to FR-12. The light only control experiments received blue light for 600 seconds (5 sec ON/25 sec OFF, 120 times) under anesthetization.

#### 2. Contextual fear conditioning test

Contextual fear conditioning task was composed of three experimental phases (habituation, fear acquisition, and fear retrieval) and conducted on two consecutive days in large sound-proof isolation chambers (Med-Associates, St. Albans, VT). On day 1, mice underwent the habituation phase for 10 min in a neutral context. The mice were then subject to the fear acquisition phase for 5 min, in which the neutral context was paired with an electric foot-shock (2 sec, 0.7 mA) delivered at 120, 180, and 240 sec, respectively. To label and subsequently manipulate a population of neurons activated during the context-shock association, blue light (5 sec) was delivered while mice receive foot-shock for 3 times.

#### 3. Sociality test

The sociality test was performed in an open field chamber (42 x 42 x 42 cm, W x D x H) as shown previously^34,35^. The test was composed of labeling session for two consecutive days and a probe test was performed 48 hours after the second labelling day. During a 5-min habituation session, test mice freely explored in the open-field chamber where two empty plastic cylinders (10 x 10 x 18 cm, W x D x H, left and right upper corner) are located at the corner. After the habituation, a novel mouse (the same sex and age matched C57B6/J mouse) was placed in one of the chambers (social chamber) and a mouse-shaped toy was placed in the other chamber (object chamber). For labeling, blue laser (5-sec duration) was delivered when the test mice enter 3-cm area around the social chamber. The laser was terminated when the test mice stayed in the area more than 5 sec. The sociality test was performed 48 hrs after the last day of labeling sessions. After the 5-min habituation, test mice explored the open-field chamber with both social and object chambers for 10 min. 589 nm yellow light was illuminated to the test mice during the next 10 min and sociality was compared. The behavior was recorded through a megapixel 720p USB camera (Shenzhen Ailipu Technology, China). The nose and both sides of the ear were tracked by DeepLabCut^36^. Staying time around each chamber was quantified as a total time when the nose or the ears were within 4-cm from the boundary of the chamber. The social preference index was measured as follows; Social preference index = Time around the social chamber/(Time around the social chamber + Time around the object chamber). The blue and yellow lasers were controlled by Pulse-Pal (Sanworks, NY, USA) and a custom-made python code.

#### 4. Seizure activity

Mice were injected i.p. with kainic acid (KA) (20 mg/kg) to induce acute seizure. This dose was used to induce generalized seizures but was sublethal^37^. After 10 min, blue light was delivered (3 sec ON/2 sec OFF; for 30 min) into the hippocampal region through the bilateral optic fibers to label neuronal population which is engaged in seizure activity. Seizure severity was classified and scored according to a modified Racine score^38–40^: 0, normal; 1, immobilization, sniffing; 2, head nodding, facial and forelimb clonus (short myoclonic jerk); 3, continuous myoclonic jerk, tail rigidity; 4, generalized limbic seizures with kangaroo posture or violent convulsion; 5, continuous generalized seizures (clonic–tonic convulsions); 6, death. The seizure only control group was not received blue light during seizure but virus injection and fiber optic implantation were made in a same way as did for the test group.

### Behavioral manipulation by optical inhibition of task-relevant neurons

Mice were allowed to recover from viral injection (pAAV-CMV-Myc-TM-KA2-CaM-NES-TEV-N-AsLOV2-TEVseq-tTA: pAAV-hSYN-M13-TEV-C-P2A-tdTomato: pAAV-TRE-eNpHR-EYFP, 1: 1: 2 ratio) and optic fiber implantation surgery. For optical inhibition of skilled motor learning, probe test (589 nm, 2 sec ON/1 sec OFF; for 45 min) was conducted on the third day after the last training session. The power of laser was measured at the end of the tip of an optic fiber and adjusted to be 15-20 mW as described previously^8^. For optical inhibition of fear memory, yellow light illumination (3 sec ON/2 sec OFF) begins when mice were subjected into the same context 24 hrs after fear acquisition. Contextual fear retrieval was measured by recording freezing responses to the conditioning context for 5 min. Freezing (%) was scored every 10 sec by two researchers blinded to the group assignments, with their scores being crossed-checked with correlation analysis. For optical inhibition of social cognition behavior, labeled social cognition-related neuronal population was inhibited by the delivery of yellow light (3 sec ON/2 sec OFF) through the bilateral optic fiber on the second day after the blue light labeling. The number of social interactions and time around chambers were analyzed as behavioral readouts. For optical inhibition of seizure responsive neurons, mice were received the second KA injection to re-induce the seizure. 589 nm yellow light was illuminated into the same hippocampal area to inhibit neural activity (by activation of eNpHR). The mean seizure score and percent of time in tonic seizure were compared between the first- and the second day KA-injected mice, and mice groups in the presence/absence of yellow light. To label hippocampal neurons that are independent of seizure, blue light was illuminated when mice were receiving foot-shock in the chamber (the same contextual fear conditioning box). Bilateral 589 nm light was delivered into the hippocampus of fear memory-labeled mice (3 sec ON/2 sec OFF) during seizure induced by the KA injection. The seizure scores were analysed.

### Tissue fixation and acquisition of confocal images

The general procedures for tissue fixation was performed as shown in a previous study^8^. Briefly, animals were deeply anesthetized by a mixture of ketamine and xylazine and then perfused transcardially, first with PBS (pH 7.4) and then with 4 % paraformaldehyde (PFA) dissolved in PBS. The brains were removed and postfixed in 4 % PFA overnight at 4 °C. The brains were embedded into 10 % melted gelatin solution for 50 min at 50°C and then gelatin solution was refreshed. Gelatin solution with the embedded brains were kept in 4 °C for ~30 min for solidification of gel. The gel was trimmed to a small cube around the brain and the cube was kept in 4 % PFA solution overnight. Coronal section (thickness, 100 μm) was made using a vibratome (Leica VT1200) for confocal imaging. Imaging was performed using upright confocal laserscanning microscope (LSM880, Zeiss, Germany) with 20×/0.8 M27 objective lens. The green to red ratio (G/R) value of individual cells were analyzed using ImageJ (NIH).

### Immunohistochemistry

After labeling by ST Cal-Light, the mice were immediately perfused with 4% paraformaldehyde in phosphate buffer solution (PBS). The brain was removed and sliced with 100 μm thickness. Slices were washed three times in PBS and blocked with 10% Normal Goat Serum in PBS with 0.1% Triton-X (Sigma) for 1 hour at room temperature (RT). Thereafter, the primary antibody was applied to the slices at 4°C for 1-2 days (1:500 for PV and CaMKII). After washing the slices three times, the slices were incubated with secondary antibodies with Alexa flour 633 or 647 (1:300, Thermo Fisher Scientific) for 2 hours at RT. To prevent bleaching of fluorescent signals, VECTASHIELD Hard set with DAPI (Vector Laboratories, Inc. Burlingame, USA) was applied to the slices on a slide glass. The images were captured using confocal microscopy (Zeiss 800, Zeiss).

### Vector design for conditional overexpression of ST-KA2 in ROSA26 Locus

Plasmid design and ST-Cal-Light knock-in mouse were generated by Ingeneous targeting laboratory (Ronkonkoma, NY). The vector was designed to have the expression of the TM-KA2-CaM-NES-TEV-N-AsLOV-TEV-tTA knock-in sequence under the pCAG promoter and also be controlled by a floxed stop cassette. The TM-KA2-CaM-NES-TEV-N-AsLOV-TEV-tTA sequence was cloned into the MluI site of the ROSA26-pCAG stop backbone vector using conventional cloning method. The stop cassette consists of a floxed PGK/gb2neoPGKpolyA2XSV40pA sequence. The knock-in sequence is followed by BGHpA sequence. The targeting vector contains a short homology arm (SA) with 1.08 kb ROSA26 genomic sequence upstream of the pCAG promoter and a 4.34 kb long homology arm (LA) downstream of BGHpA sequence. The targeting vector was confirmed by restriction enzyme analysis and sequencing after each modification. The boundaries of the pCAG-stop cassette and BGHpA-ROSA26 genomic sequences were confirmed by sequencing with primers ROSASQ1 and ROSASQ2.

In order to test the efficacy for labeling of active subset of neurons using newly generated Cal-KI mice *in vivo* (Fig. 5), pre-mixed viral solution (pAAV-M13-TEV-C-IRES-SP6-Cre: pAAV-TRE-EGFP, 1:4) was injected to the M1 (Coordinates: AP, +0.25 mm; ML, +1.5 mm from bregma, and DV −0.2 mm from the brain surface; see Fig. 5d). The optic fiber implantation coordinates for the labeling of M1 neurons in ST-Cal-Light KI mice is described above (see stereotaxic surgeries section).

### Generation and verification of double transgenic mice

Male and female ST-Cal-Light KIs and WT littermates were maintained on C57BL/6 background. For identification of active PV interneurons, KI heterozygous (Cal-Het) mice were bred with Cal-Het::PV-Cre mice to generate Cal-Het (or Hom) and WT littermates that were PV-Cre hemizygous (see Fig. 5i). PV-Cre mice were derived from a mouse knock-in of Cre recombinase directed by the PV promotor/enhancer (Pvalbtm1(cre)Arbr, The Jackson Laboratory; Stock No: 017320). The same procedure for the generation of Cal-Het::EMX-Cre was used with the following verification. For targeting expression into either M1 pyramidal or PV neurons, 700 nl viral stock solution (pAAV-hSYN-M13-TEV-C-P2A-tdTomato: AAV1-CMV-PI-Cre-rBG: pAAV-TRE-EGFP = 1: 2: 4 ratio) was injected to the same target site for the manipulation of the labeled subset of neurons in Cal-Het::EMX-Cre or Cal-Het::PV-Cre mice, respectively (see above fiber implant coordination).

### Pharmacology

Bicuculline and TTX were purchased from Tocris Bioscience (Minneapolis, MN, USA). All drug stock solution was prepared with 1,000x or greater.

### Statistics

Statistical significance of culture neuron data was calculated by one-way ANOVA with post hoc Bonferroni test using SPSS 21 (IBM) software. Organotypic slice culture and *in vivo* data ware analyzed using IgorPro (version 6.10A; WaveMetrics, Lake Oswego, OR, USA). Statistical data were presented as means ± standard error of mean (s.e.m., denoted as error bars), and *n* indicates the number of cells or animals studied. The significance of differences between two experimental conditions was evaluated using Student’s *t* test, or Wilcoxon’s signed rank test for non-paired and paired data after testing normality using the Kolmogorov-Smirnov test. Comparison of G/R in *in vivo* and *in vitro* slice culture experiments and behavioral experiments was evaluated using nonparametric Mann-Whitney *U* test. Difference were considered as significant when p < 0.05. n.s.: no statistical significance; *: p < 0.05, **: p < 0.01, ***: p < 0.005.

